# SNP-ChIP: A versatile and tag-free method to quantify changes in protein binding across the genome

**DOI:** 10.1101/390005

**Authors:** Luis A. Vale-Silva, Tovah E. Markowitz, Andreas Hochwagen

**Affiliations:** Department of Biology, New York University, New York, NY 10003, USA

## Abstract

Chromatin-immunoprecipitation followed by sequencing (ChIP-seq) is the method of choice for mapping genome-wide binding of chromatin-associated factors. However, broadly applicable methods for between-sample comparisons are lacking. Here, we introduce SNP-ChIP, a method that leverages small-scale intra-species polymorphisms, mainly SNPs, for quantitative spike-in normalization of ChIP-seq results. Sourcing spike-in material from the same species ensures antibody cross-reactivity and physiological coherence, thereby eliminating two central limitations of traditional spike-in approaches. We show that SNP-ChIP is robust to changes in sequencing depth and spike-in proportions, and reliably identifies changes in overall protein levels, irrespective of changes in binding distribution. Application of SNP-ChIP to test cases from budding yeast meiosis allowed discovery of novel regulators of the chromosomal protein Red1 and quantitative analysis of the DNA-damage associated histone modification *γ*-H2AX. SNP-ChIP is fully compatible with the intra-species diversity of humans and most model organisms and thus offers a general method for normalizing ChIP-seq results.

## Introduction

Chromatin immunoprecipitation followed by DNA sequencing (ChIP-seq) has emerged as the method of choice for mapping the genome-wide distribution of proteins and protein modifications and has led to important discoveries in both basic chromatin biology and disease states ^1,2^. A core result of ChIP-seq experiments is the generation of genome-wide target signal tracks, which are obtained from read pileups, typically normalized against a mock, non-immunoprecipitated control sample (input sample). Signal tracks are used for identification of regions with elevated numbers of mapped reads (peaks) as well as other downstream analyses ^3^. However, because of the necessary internal normalization procedures, signal tracks can only be used for comparisons between samples if a method for inter-sample normalization is available ^3^. This is a crucial, often overlooked, caveat of ChIP-seq, as well as other genome-wide biochemical analysis methods relying on next-generation sequencing ^4^

For sparsely bound proteins, such as transcription factors, inter-sample normalization can often be achieved using statistical methods ^2^ or ChIP followed by real-time quantitative PCR (ChIP-qPCR) ^5^. These methods, however, either assume constant global signal or a constant signal at selected genes as basis for normalization, which is difficult to verify, in particular for more broadly distributed factors. The solution to overcome this limitation is the addition of a “spike-in” reference sample ^2,6^. The spike-in procedure consists of adding a constant amount of exogenous material to all tested samples, ideally prior to any critical steps in the experimental protocol. Provided that the spike-in material contains a target that is bound by the antibody as efficiently as the study target and that the resulting sequencing reads can be distinguished from the test sample, the number of spike-in reads should be the same across all tested samples. The spike-in thus functions as an internal control against which to normalize the ChIP-seq results ^6^. Spike-ins are well established for RNA-seq analyses where use of RNA from a different species allows simple sequence-based distinction between test sample and spike-in ^7^. The additional requirement for cross-reactivity of the antibody in ChIP-seq experiments, however, effectively restricts the applicability of inter-species spike-ins to a limited set of highly conserved proteins ^8,9^.

Ways to broaden the applicability of ChIP spike-ins include either tagging proteins in the test and spike-in samples with a common epitope ^10^, or using a second, spike-in specific antibody against a natural ^11^ or a synthetic target ^12^. These strategies, however, come with their own specific drawbacks. The use of protein tagging adds the potential for prohibitive disruption of protein function and is incompatible with the analysis of protein modifications. The use of a second, spike-in specific antibody, on the other hand, requires labor-intensive technical validation of the compatibility of the second antibody and no longer controls for biases in the immunoprecipitation step between samples.

Here, we show that these issues can largely be overcome by using spike-in material from the same species. This approach, which we name SNP-ChIP, enables reproducible semi-quantitative measurement of global protein levels and also works for protein modifications and fast evolving proteins.

## Results

### Experimental rationale of SNP-ChIP

The basic premise of SNP-ChIP is that cells from the same species can serve as spike-in material provided it harbors sufficient genetic diversity, primarily in the form of single-nucleotide polymorphisms (SNPs). The signal at each polymorphism provides an independent measure of test-sample/spike-in ratio that together allows calculation of a normalization factor and appropriate scaling of ChIP-seq results (Fig. 1a). If there is sufficient genetic diversity to allow a large fraction of sequencing reads to be assigned to the genomes of origin, SNP-ChIP additionally allows the generation of genome-wide target distribution profiles. Importantly, because SNP-ChIP uses the same species as the source of the spike-in material, it will work with virtually any target in the organism’s proteome, including post-translational modifications, provided a ChIP-grade antibody is available.

**Figure 1.**
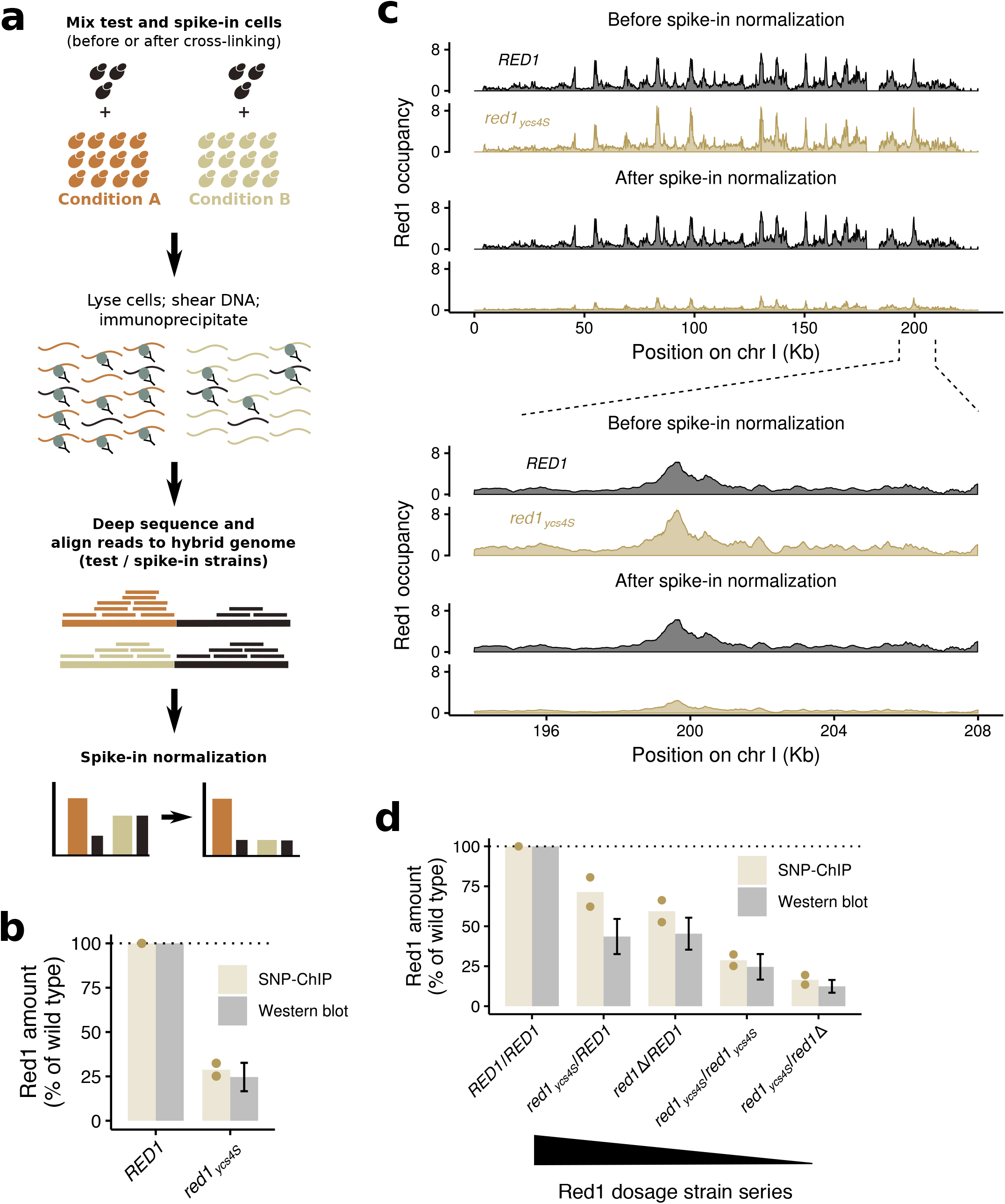
SNP-ChIP adds the ability to measure semi-quantitative amounts of target protein to traditional ChIP-seq. (a) Main steps of SNP-ChIP exemplified for two hypothetical conditions. (b) Target protein Red1 levels produced by SNP-ChIP (equivalent to the wild type-normalized spike-in normalization factor), compared to previously published levels measured by western blot (mean +/− S.E.M.) ^19^. Points represent individual replicate values and bars represent average value. (c) Fragment pileup produced using MACS2 with SPMR sequencing depth normalization (fragment pileup per million reads) of an example chromosome and chromosome region before (top panel) and after (bottom panel) spike-in normalization. (d) Target protein Red1 levels for Red1 dosage strain series compared to previously published levels measured by western blot (mean +/− S.E.M.) ^19^, as in (b).

### SNP-ChIP of a rapidly evolving chromosomal protein

To test the power of intra-species spike-ins, we focused on yeast meiotic recombination, which involves many broadly distributed chromosomal proteins and post-translational modifications. One typical example is the axial-element protein Red1, which plays important roles in meiotic recombination. Red1 is broadly bound along chromosomes ^13–15^ but, like other meiotic factors, its sequence has diverged even in closely related species ^16^. Furthermore, like many proteins, Red1 cannot easily be tagged without disrupting protein function ^15,17^. These attributes mean Red1 is not amenable to standard spike-in approaches, making it a particularly suitable target for SNP-ChIP. Moreover, mutations that change the overall levels and chromosomal distribution of Red1 are available ^15,18,19^, providing benchmarks for evaluating the efficacy of SNP-ChIP.

SNP-ChIP of Red1 was performed using the SK1 genetic background ^20^ as test strain and a meiosis-optimized variant of the widely used S288c reference strain as spike-in ^21,22^. For both genetic backgrounds, high-quality end-to-end genome assemblies are available ^23^. These assemblies differ by about 76,000 single-nucleotide polymorphisms (SNPs), spaced at a median distance of 70 bp (Fig. S1), which constitutes enough variation to allow unambiguous assignment of a large proportion of sequencing reads. To perform SNP-ChIP, test cells (SK1) were mixed with a constant fraction of meiotic spike-in cells (S288c) before subjecting the mixtures to a standard ChIP-seq protocol. The generated reads were aligned to a hybrid genome built by concatenating genome assemblies of the test and spike-in genomes. Reads were aligned with perfect match conditions, excluding any reads aligning to more than one location. Consequently, any reads overlapping at least one SNP were assigned to a specific genome and genomic location, while reads not overlapping a polymorphism mapped to both genomes and were thus discarded.

We initially investigated the ability of SNP-ChIP to detect changes in chromatin association resulting from reduced protein production. The *red1_ycs4S_* allele is caused by a mutation in the promoter of *RED1* that leads to a reduction of Red1 levels to about 20-25% of wild type and a near complete loss of cytologically observable axial elements ^19^. Importantly, traditional ChIP-seq analysis was unable to detect this change in protein abundance and produced indistinguishable Red1 profiles between wild type and *red1_ycs4S_* mutants ^19^. By contrast, when we applied SNP-ChIP to compare these two strains, the reduced Red1 binding levels were readily apparent (Fig. 1b). Calculation of a spike-in normalization factor based on the relative abundance of total sample and spike-in reads yielded a Red1 level in the *red1_ycs4S_* mutant of 28.8 ± 5.1% (S.D.) of the wild type, closely matching the reported change in Red1 levels obtained from western analysis ^19^. This normalization factor allowed appropriate signal scaling of ChIP-seq profiles for the two conditions (Fig 1c).

SNP-ChIP was further validated by applying it to a Red1 dosage series, which consists of different combinations of *RED1* alleles (*RED1, red1_ycs4S_, red1*Δ) yielding a stepwise decrease in Red1 levels (Fig. 1d) ^19^. SNP-ChIP measurements of Red1 chromatin association in this series again closely matched previously published protein levels (Fig. 1d). In fact, SNP-ChIP measurements appeared more accurate than quantitative western analysis, which failed to resolve the expected reduction in protein levels between *RED1/red1_ycs4S_* and *RED1/red1*Δ. cells ^19^. Taken together, these data show that SNP-ChIP accurately measures reductions in global Red1 binding over a wide range of target protein levels.

### SNP-ChIP is robust to variation in sequencing depth and fraction of spike-in cells

We used several approaches to probe the technical robustness of SNP-ChIP. High-throughput sequencing technologies produce variable numbers of reads per sample, depending on factors like sequencing instrument and sample mutiplexing. To model the effect of lower sequencing depth on the reproducibility of SNP-ChIP analysis, we subsampled the reads of the immunoprecipitated and input samples from wild type and *red1_ycs4S_* test conditions to different depths (ranging from 1 to 10 million reads). Plotting subsample size against number of aligned reads showed a perfectly linear correlation for all samples (Fig. 2a), indicating that a wide range of sequencing depths will yield robust quantitative information by SNP-ChIP. We computed the spike-in normalization factor using all 10,000 possible combinations of read subsamples (10 read subsamples for each of four sequenced samples) and found a very tight distribution of results (0.2848 ± 0.0015, S.D.; Fig. 2b). This establishes that sequencing depth does not need to be balanced between immunoprecipitated and input samples, or between different conditions, to produce accurate proportions of reads mapping to the test and the spike-in genomes.

**Figure 2.**
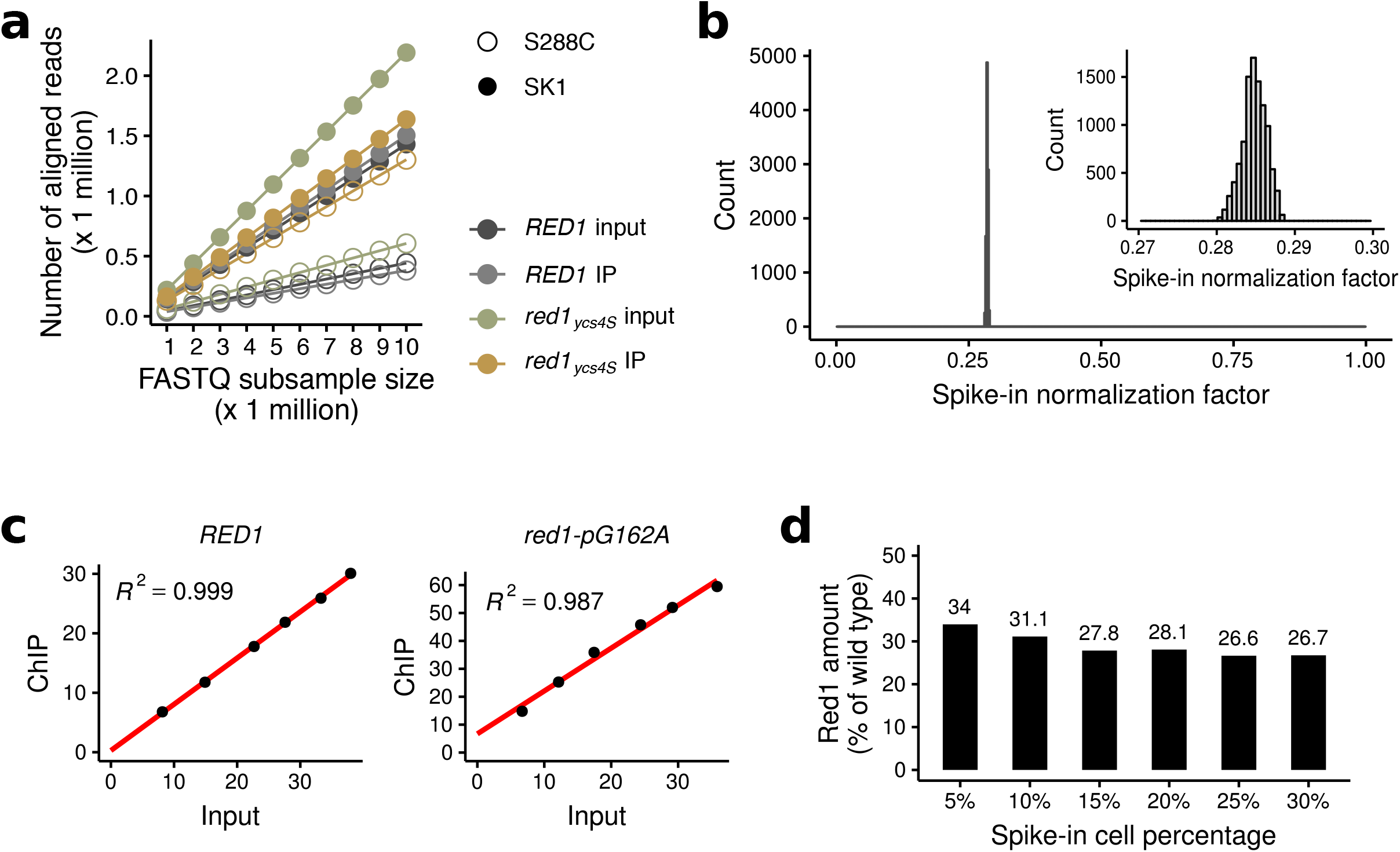
SNP-ChIP is robust to variation in both sequencing depth and amount of spike-in cells. (a) Number of aligned reads in SNP-ChIP as a function of total size of raw read sample. Systematic raw read subsamples of size 1 to 10 million were obtained for each sample and mapped to the hybrid SK1-S288c genome. (b) Distribution of spike-in normalization factors computed using all 10’000 possible combinations of read subsamples in (a). The inset is a zoom-in on a narrow window of the x axis where all values are located. (c) Proportion of reads aligning to the spike-in genome in the input sample versus the immunoprecipitated sample (ChIP) for wild type and *red1-pG162A* strains. Note: In contrast to *red1_ycs4S_*, which contains an introgressed region with dozens of SNP surrounding the *RED1* locus, the *red1-pG162A* mutant only carries the causative promoter mutation. (d) Resulting wild type-normalized spike-in normalization factor (equivalent to Red1 amount) in the *red1-pG162A* strain after performing SNP-ChIP with percentages of added spike-in cells ranging from 5 to 30%.

Another condition that may affect the results of SNP-ChIP is the amount of spike-in material added to the samples. Spike-in normalization methods assume a linear relationship between the amount of spike-in material and the resulting proportion of spike-in reads in the immunoprecipitated sample. This condition is essential for the results to be independent from the amount of spike-in material. To verify this assumption, we prepared samples with spike-in cell proportions ranging from 5 to 30 percent. As test samples we used wild type and a strain with a single *red1-pG162A* promoter mutation that phenocopies the *red1_ycs4S_* allele ^19^. As shown in Fig. 2c, the proportion of spike-in reads in the input samples (reflecting the amount of spike-in material added to the test sample) correlates linearly with the resulting proportion of spike-in reads in the immunoprecipitated sample, for both the wild type and the *red1-pG162A* sample. Moreover, the *red1-pG162A* sample yielded a very similar normalization factor to the *red1_ycs4S_* allele, further supporting the robustness of the method. Low spike-in cell percentages (5 and 10%) resulted in somewhat increased estimates of the normalization factor (Fig. 2d), however, likely due to increased noise. These results suggest that spike-in material proportions of 15% and higher are appropriate for SNP-ChIP. All other experiments shown here used a spike-in proportion of 20%.

Finally, we investigated the impact of the calculation method to compute the spike-in normalization factor. The SNP-ChIP normalization factor calculated in the examples shown so far relies on total read counts aligned to the test and the spike-in genomes (see Methods section). An alternative method is to compute the scalar mean value of the aligned read pileup score. We tested the utility of this alternative by calculating the pileup score at (1) all genomic positions, (2) at SNP positions only, or (3) at SNP positions falling within called signal peaks. The last approach will effectively exclude regions expected to hold only background signal, along with any false negative regions. We found very similar values and high concordance between all four methods in all cases (Fig. S2a), although read pileups consistently produce slightly lower values than the read count method (Fig. S2b). Overall, however, the difference is relatively small and we believe the read count-based method, which is computationally much simpler, represents an appropriate approximation, at least for broadly distributed proteins.

### Binding profiles obtained directly from SNP-ChIP experiments

The primary utility of SNP-ChIP is the generation of a normalization factor that allows scaling of profiles obtained by traditional ChIP-seq experiments run under the same conditions (Fig. 1c). Given the broad distribution of SNPs across the two analyzed genomes, we explored the possibility that SNP-ChIP could also directly yield informative binding profiles, even though this application is clearly limited by the available SNP density. Comparing a sample sequenced with spike-in to data obtained using a replicate, non-spiked sample ^15^ shows that signal tracks of spiked samples closely mirror those of the non-spiked control, although some signal gaps can be seen in the spiked sample (Fig. S3a), in particular at higher resolution (indicated by the red arrow). Thus, as expected, the use of same-species spike-in causes some loss of information. This issue appears negligible for broad peaks, as called peaks show a very close agreement (Fig. S3b). Narrow peaks show more disagreement, with only about one third of the called peaks overlapping between the two samples. These data indicate that SNP-ChIP can also provide direct information about protein distribution, in particular for larger-scale binding patterns.

### Global Red1 levels are reduced in cohesin and *hop1*Δ mutants

We sought to apply SNP-ChIP to investigate mutant situations that cause a broad protein redistribution. Redistribution is a challenge for traditional quantification methods, such as ChIP-qPCR, because identifying regions that remain unbound is non-trivial. In the absence of conserved cohesin subunit Rec8, Red1 distribution along the genome changes dramatically, displaying large regions of depletion alternating with dense clusters of binding ^15,18^. Whether overall binding levels of Red1 change in *rec8*Δ mutants, however, remains unclear. We employed SNP-ChIP to address this question and found a pronounced decrease of overall Red1 binding levels (Fig. 3a). Direct comparison of Red1 occupancy along two example chromosomes illustrates both the dramatic redistribution and the overall decrease in Red1 binding compared to wild type (Fig. 3b). Thus, Rec8-cohesin is essential for the full chromosomal enrichment of Red1.

**Figure 3.**
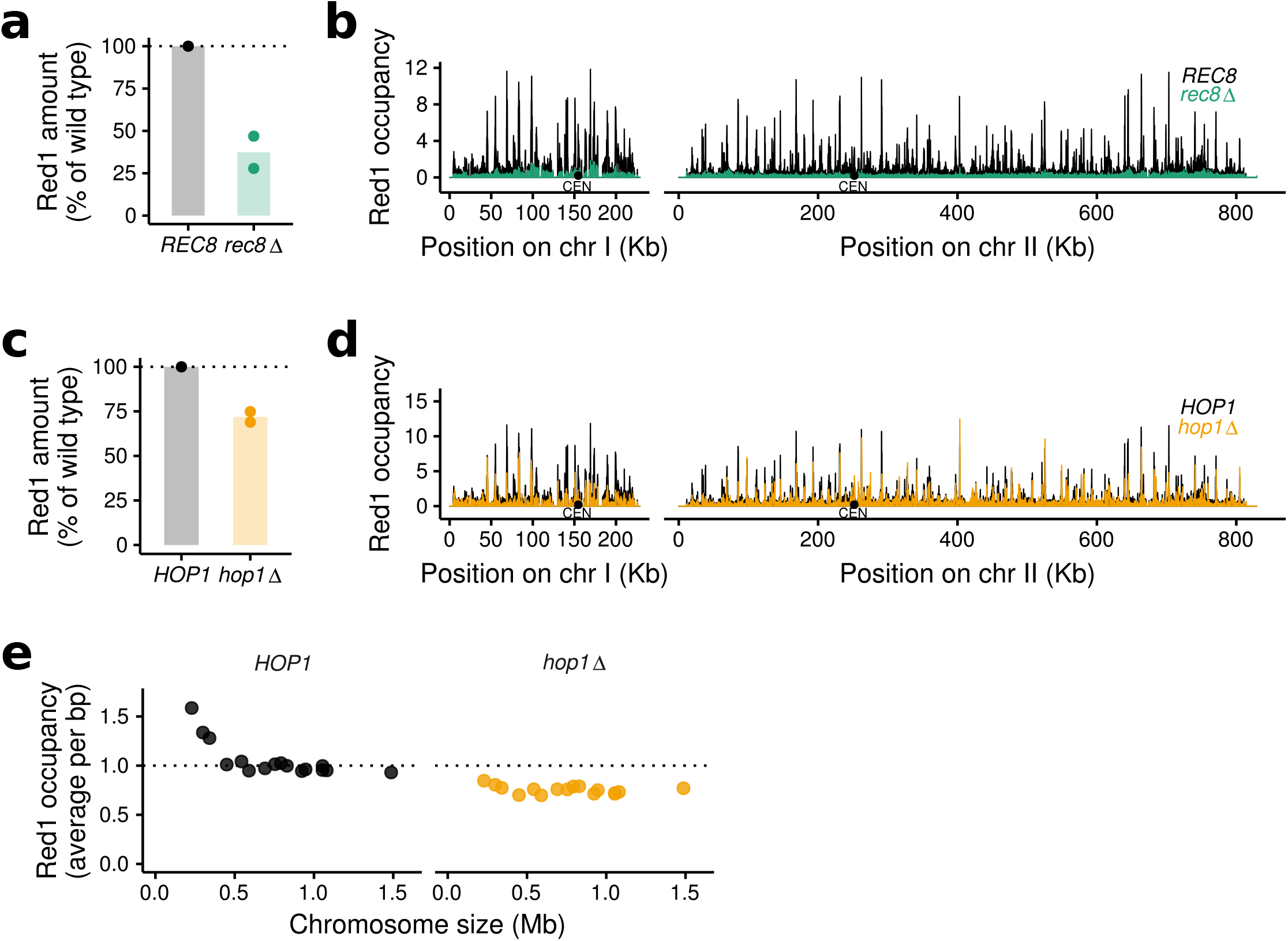
SNP-ChIP analysis in mutants with large-scale target redistribution. (a) Target protein Red1 levels in *rec8*Δ mutant relative to wild type produced by SNP-ChIP. Points represent individual replicate values and bars represent average value. (b) Spike-in-normalized fragment pileups in wild type and *rec8*Δ mutant strains produced using MACS2 with SPMR sequencing depth normalization (fragment pileup per million reads) plotted on two example chromosomes. (C) Red1 levels in *hop1*Δ mutant relative to wild type produced by SNP-ChIP (as in a). (d) Spike-in-normalized fragment pileups in wild type and *hop1*Δ mutant strains (as in b). (e) Spike-in-normalized average Red1 signal on individual chromosomes in *hop1*Δ mutant.

Hop1 is another important protein of the yeast axial element ^24^ that physically interacts with Red1 ^17^. Axial-element proteins are recruited in higher amounts to small chromosomes, but in the absence of Hop1, Red1 binding becomes less dependent on chromosome size ^15^. Previous work using *in silico* scaling (NCIS) ^25^, suggested that this reduction resulted from a selective increase in Red1 recruitment to large chromosomes ^15^. NCIS, however, requires the definition of genomic regions that are unbound, which is difficult to ascertain with broadly distributed chromosomal proteins like Red1. Therefore, we reinvestigated this question by performing SNP-ChIP of Red1 in a *hop1*Δ mutant. SNP-ChIP reproduced the previously found weakening of chromosome-size bias. However, the spike-in normalization factor showed an overall decrease of Red1 recruitment to 71.9 ± 4.2% of the wild type Red1 amount (Fig. 3c, d). This decrease is particularly strong on small chromosomes (Fig. 3e). We note that mild loss of Red1 binding does not generally result in a loss of chromosome-size bias, because deletion of the histone methyltransferases Set1 and Dot1 causes similar ~ 20% reductions of overall Red1 recruitment levels but does not affect the distribution of Red1 binding among chromosomes (Fig. S4). These data suggest that loss of Hop1 leads to a generalized reduction of Red1 signal across all chromosomes that particularly affects the three smallest chromosomes.

### *γ*-H2AX levels do not change in Red1 dosage strain series

To test if SNP-ChIP also allows quantitative analyses of protein modifications, we targeted phosphorylation of histone H2A on serine 129 (*γ*-H2AX). This modification is rapidly induced following the formation of DNA double-strand breaks (DSBs) ^26^. In mitotic yeast, the *γ*-H2AX modification spreads about 50 kb on either side of a DSB ^27,28^. In addition, constitutive *γ*-H2AX is found near telomeres throughout the cell cycle ^29^. To analyze the distribution and DSB dependence of *γ*-H2AX in meiosis, we performed SNP-ChIP in a wild type strain, as well as the Red1 dosage series, which shows a mild (up to 30%) reduction in DSB levels^19^, and a *spo11-Y135F* mutant, encoding a catalytically dead Spo11 protein that does not form meiotic DSBs ^30,31^.

Measuring *γ*-H2AX levels in meiosis revealed no difference between the wild type and any of strains with reduced Red1 levels, irrespective of calculation method (Fig. 4a, b). The uniform signal along chromosomes is consistent with the spreading of the *γ*-H2AX mark from all yeast DSB hotspots, which are distributed throughout the whole genome, and likely explains why a mild reduction in DSB levels does not lead to a noticeable drop in global *γ*-H2AX signal. The *spo11-Y135F* control, on the other hand, displayed only about 25% of the wild type *γ*-H2AX levels. Signal was markedly enriched next to telomeres, with the interstitial regions only showing weak signals likely associated with gene expression ^32^. These data show that the constitutive telomere-associated *γ*-H2AX signal is also maintained in meiotic prophase. Moreover, scaling of signal tracks indicates that telomere-adjacent *γ*-H2AX signal remains largely unchanged in the *spo11-Y135F* mutant, consistent with the fact that meiotic DSB formation is nearly undetectable in these regions ^33^. Together, these data show that SNP-ChIP allows quantitative comparisons between ChIP-seq experiments regardless of the antigen and thus provides a versatile and tag-free method for measuring global chromatin associations.

**Figure 4.**
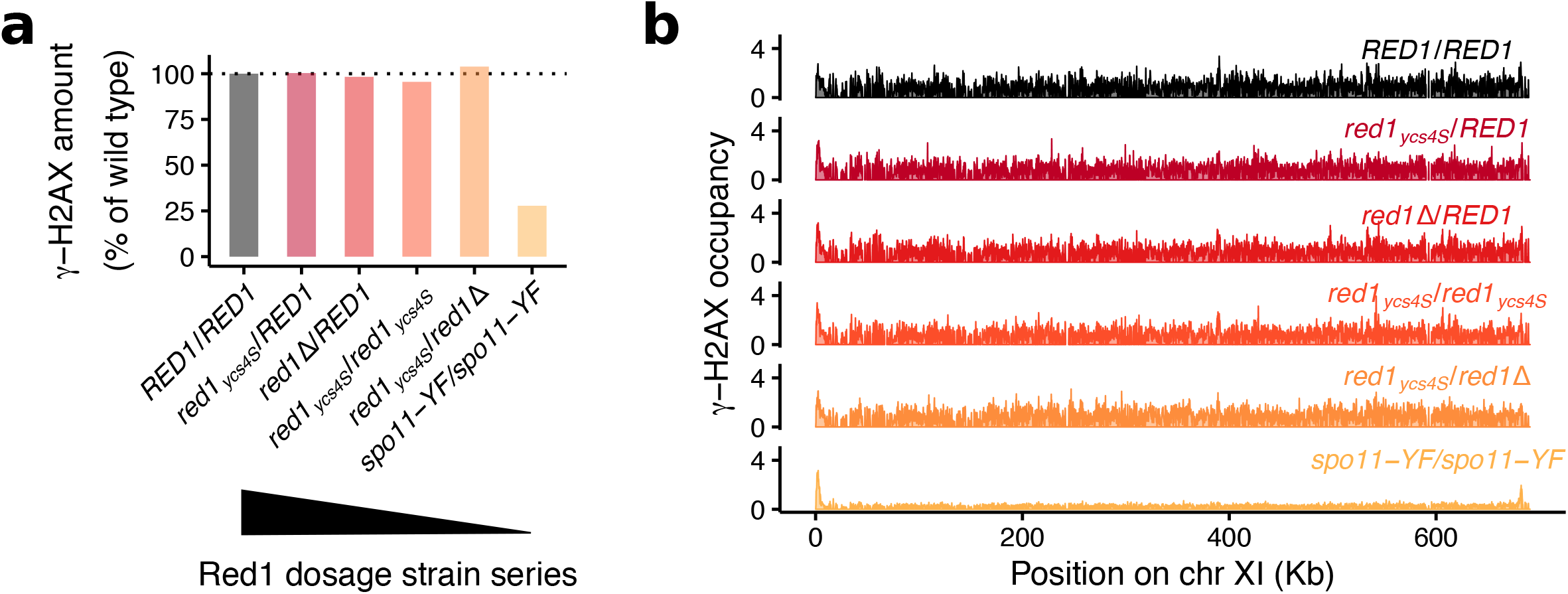
SNP-ChIP analysis of a protein modification. (a) *γ*-H2AX levels for the Red1 dosage strain series relative to wild type produced by SNP-ChIP. (b) Fragment pileup produced using MACS2 with SPMR sequencing depth normalization (fragment pileup per million reads) on an example chromosome after spike-in normalization.

## Conclusion

Our data show that small-scale intra-species genetic polymorphisms can be leveraged for quantitative spike-in normalization of ChIP-seq results. Sourcing spike-in material from the same species largely preserves antibody cross-reactivity and thus will work with virtually any target in an organism’s proteome without the need for epitope tagging. It also ensures complete physiological coherence between the test and the spike-in cells, thereby avoiding biases at experimental steps such as chromatin fixation or cell lysis.

The primary output of SNP-ChIP is a normalization factor that can be used to appropriately scale ChIP-seq profiles. Because the normalization factor relies on combined measurements of thousands of SNPs it is highly robust to variations in sequencing depth or changes in protein distribution between samples. In multiplexed libraries, SNP-ChIP can therefore be performed with relatively low sequencing coverage alongside traditional ChIP-seq experiments to yield the necessary scaling information.

SNP-ChIP can also provide substantial positional information, although this application is necessarily limited by the availability of high-confidence SNPs. Our experiments using yeast strains with ~0.7% sequence divergence and 100-nt long reads showed that the method generated sufficient resolution to recover genomic regions of Red1 enrichment. Moreover, preliminary experiments indicate that using longer reads further minimizes gaps (data not shown). Thus, SNP-ChIP can provide high-quality pilot information for subsequent ChIP-seq analyses at higher read depth.

The reliance on thousands of SNPs also means that SNP-ChIP will be particularly powerful for the quantitative analysis of broadly distributed proteins and chromatin marks. Applying SNP-ChIP to proteins that interact with chromatin in more specific, highly localized positions (e.g. transcription factors), will likely result in a disproportionate number of SNPs exhibiting background signal that will affect the calculation of the normalization factor. Indeed, preliminary experiments testing the budding yeast transcription factor Gal4 suggested that SNP-ChIP is not ready to handle such targets (data not shown). While SNP-ChIP generated reliable signal track data, the normalization factor computation method does not work as-is and will require further development. We note, however, that sparsely binding proteins are inherently more tractable targets for ChIP-qPCR, thus reducing the need for a spike-in method.

SNP-ChIP is fully compatible with the intra-species genetic diversity of humans and most model organisms ^34^ and should be applicable to any experimental system for which a reliable collection of high-quality SNPs is available. In preliminary *in silico* experiments testing decreasing numbers of SNPs, the method generated stable normalization factors with as low as 0.01% sequence divergence (equivalent to about 1,200 SNPs in the yeast genome; data not shown). Thus, we expect that SNP-ChIP will allow semi-quantitative mapping of a wide range of chromatin binding factors and modifications that have so far stood beyond the reach of quantitative ChIP-seq methods.

## Methods

### Strains and meiotic time courses

All strains used are listed in Table S1. The test-sample strains were of the SK1 background. The spike-in material used a meiosis-optimized S288c strain that carries three SK1-derived SNPs, which improve sporulation efficiency and meiotic synchrony of S288c ^22^. To further improve synchrony of the spike-in strain, auxotropic markers were restored using plasmid insertions or PCR-based allele transfer. To induce meiosis, cells were pregrown in YPD for 24 hours at room temperature, followed by inoculation in BYTA media at O.D._600_=0.3 and growth for 16.5 hours at 30°C ^35^. Cells were washed twice with water and inoculated at O.D._600_=1.9 in 0.3% potassium acetate (pH 7.0) to induce meiotic entry. Synchronous entry was confirmed by taking hourly samples for flow cytometry analysis of DNA content.

### SNP-ChIP sample preparation

Samples were collected at 3 hours for SK1 strains or at 6 hours for the slower sporulating S288c spike-in sample. Cells were fixed in 1% formaldehyde for 30 minutes at room temperature and quenched by addition of glycine to a final concentration of 125 mM. For the experiments shown here, we fixed the spike-in cells in advance as a batch and kept frozen aliquots at −80°C. However, spike-in cells can also be prepared simultaneously with the sample cells. The number of cells in each sample was determined by counting on a hemocytometer. Unless indicated otherwise, cells from the test sample (SK1) were mixed with cells from the spike-in sample (S288c) at a ratio of 80:20 before cell lysis and ChIP.

### Chromatin immunoprecipitation (ChIP) and Illumina sequencing

ChIP was performed as described previously ^36^. Samples were immunoprecipitated with 2 μl anti-Red1 serum (Lot#16440, kind gift of N. Hollingsworth) or 2 μl anti-phospho-H2A-S129 antibody (Abcam #ab15083) per sample. Library preparation was performed as described ^15^. Library quality was confirmed by Qubit HS assay and 2200 Tape Station. 100-bp single-end sequencing was performed on an Illumina NextSeq 500 instrument.

### Read alignment

The generated reads were aligned to a hybrid genome built by concatenating high-quality genome assemblies of the test and spike-in reference genomes (SK1 and S288c) ^23^. Reads were aligned with perfect match conditions and excluding any reads aligning to more than one location. Normalization of read density was completed as described ^36^. Where indicated, peaks of enrichment were called using MACS-2.1.0. Plots show an average of two replicates. To evaluate coverage relative to standard ChIP-seq profiles, we compared SNP-ChIP results to published datasets GSE69232 ^15^ and GSE87060 ^19^.

### Calculation of the spike-in normalization factor

Let *C_endo_* and *C_spike_* be the count of all reads aligned to the endogenous and the spike-in genomes, respectively, in a given sequenced sample. Quotient *Q* is defined as the ratio between the read counts:

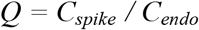

The non-immunoprecipitated, input sample’s *Q* value (*Q_Input_*) provides a measure of the percentage of the sample that is comprised of spike-in cells in the respective test condition. This consists of the experimental percentage of spike-in material actually added to the sample, corrected for any technical variation or imprecision. The ChIP sample’s *Q* value (*Q_ChIP_*) depends on the amount of target protein present in that experimental condition. A normalization factor *Nf* can thus be obtained for each condition as the ratio between the two *Q* values:

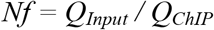

To obtain spike-in-normalized conditions, each condition is multiplied by the respective normalization factor value *Nf*. The extent to which *Q_ChIP_* differs from *Q_Input_* in each experimental condition is determined by the amount of target protein and how much that differs from the amount of target protein in the spike-in. Since the latter is constant across all tested conditions, the result of the normalization is a semi-quantitative measure of the target protein amounts, yielding normalized conditions that can be compared directly to each other.

### Data availability and codes

Data sets have been deposited in NCBI’s Gene Expression Omnibus and are accessible through GEO Series accession number GSE115092. Code used for data analysis and producing figures is available on Github (https://github.com/hochwagenlab/SNP-ChIP).

## Acknowledgments

We thank F. Winston for sharing strains and N. Hollingsworth for sharing antibodies. We are grateful to S. Ercan and S. Keeney for helpful discussions and the NYU Department of Biology Sequencing Core for technical assistance and debarcoding. This work was supported in part by NIH grant R01 GM111715 and research grant FY16-208 from the March of Dimes Foundation to A.H.

## Author contributions

L.A.V.-S. and A.H. conceived of the study. T.E.M produced key resources. L.A.V.-S. conducted the experiments. L.A.V.-S. and A.H. analyzed the data and wrote the manuscript with input from T.E.M.

